# Analytical and computational solution for the estimation of SNP-heritability in biobank-scale and distributed datasets

**DOI:** 10.1101/2024.09.20.614017

**Authors:** Guo-An Qi, Qi-Xin Zhang, Jingyu Kang, Tianyuan Li, Xiyun Xu, Zhe Zhang, Zhe Fan, Siyang Liu, Guo-Bo Chen

## Abstract

Estimation of heritability has been a routine in statistical genetics, in particular with the increasing sample size such as biobank-scale data and distributed datasets, the latter of which has increasing concerns of privacy. Recently a randomized Haseman-Elston regression (RHE-reg) has been proposed to estimate SNP-heritability, and given sufficient iteration (*B*) RHE-reg can tackle biobank-scale data, such as UK Biobank (UKB), very efficiently. In this study, we present an analytical solution that balances iteration *B* and RHE-reg estimation, which resolves the convergence of the proposed RHE-reg in high precision. We applied the method for 81 UKB quantitative traits and estimated their SNP-heritability and test statistics precisely. Furthermore, we extended RHE-reg into distributed datasets and demonstrated their utility in real data application and simulated data. The software for estimating SNP-heritability for biobank-scale data is released: https://github.com/gc5k/gear2.

## Introduction

Estimating heritability has been one of the central tasks in statistical genetics (Visscher *et al*., 2008). Given the increasing sequencing capability, high-throughput genetic data have been emerging in the form of biobank-scale (Bycroft *et al*., 2018), which challenges statistical computation, in particular, such as the estimation of heritability for complex traits. Conventional methods for estimating heritability, like the linear mixed model, often takes computational cost of 𝒪(*n*^2^*m* + *n*^3^), where *n* is the sample size and *m* is the number of markers. These costs can become infeasible in the context of biobank-scale data. Haseman-Elston regression (HE) was originally proposed for linkage analysis (Haseman and Elston, 1972). After the nuclear correlation between sib pairs is replaced by the linkage disequilibrium (LD) for unrelated samples, the modified HE can be used to estimate heritability and is much faster than REML (Chen, 2014). Given that *m* is often greater than *n* given the current data, any calculation that is upon genetic relationship matrix (GRM) will be unfavorable even for HE. To reduce the computational cost of estimating heritability, recently a randomized estimation of heritability has been introduced (Wu and Sankararaman, 2018), called randomized Haseman-Elston regression (RHE-reg), which is a promising method that can be used for both single-trait and bi-trait analyses (Wu *et al*., 2022).

RHE-reg is built on a hybrid framework, which has favoured analytical properties of the Haseman-Elston regression and the feasible computational cost of 𝒪(*nmB*) for biobank-scale data. As pointed out by a recent systematic review, iteration control poses one of the challenges for RHE-reg (Tang *et al*., 2022). However, the original report by Wu and Sankararaman did not give a clear solution for the round of iteration (*B*) (Wu and Sankararaman, 2018). In this study, we investigated RHE-reg and found an analytical procedure to control *B*, which can provide customized iteration for a given data.

Now, one of the trends is that genomic cohorts are mushroomed such as emerging non-invasive prenatal testing cohorts (Yang *et al*., 2024; Xiao *et al*., 2023), but the bottleneck is how to share genomic data without compromising personal privacy (Chen *et al*., 2024). As recently practiced, when genotypes have been masked in randomization, the randomized method has proven to be reliable in addressing genetic problems for distributed data, such as searching relatives (Zhang *et al*., 2024; Chen *et al*., 2017). Following this idea, it is found that after the randomization step, RHE-reg can be modified to estimate heritability for distributed datasets, reminiscent of vertical or horizontal federated learning (McMahan *et al*., 2017).

## Method description

Wu and Sankararaman proposed a randomized implementation for the Haseman-Elston regression (RHE-reg), which dramatically reduced the computational time from 𝒪(*n*^2^*m*) to 𝒪(*nmB*) in dealing with *tr*(***K***^2^) (Wu and Sankararaman, 2018). It is clear that a large *B* is helpful in improving precision, but it is unsolved how to get a estimate for *B* and its role in determining the boundaries of key statistics (Tang *et al*., 2022). This work is in general consistent with Wu and Sankararaman’s work, but we present the analytical sampling variance of the estimated *h*^2^ and its corresponding test statistics. We can consequently evaluate how *B* influence the estimation of heritability and its corresponding *z* score, and, as data can be very large, the control of *B* is of theoretical as well as practical importance. An analytical resolution crystalizes a computational procedure, and we further extend the method to another two new scenarios, called vertical-RHE-reg, which is a global implementation for LD score regression (Bulik-Sullivan *et al*., 2015), and horizontal-RHE-reg, which enables Federated Learning but we estimate heritability in distributed data without compromise of privacy (McMahan *et al*., 2017).

## Materials and Methods

### A framework for Randomized Haseman-Elston regression (RHE-reg)

In essence, Haseman-Elston regression is a kind of method of moments (MoM) estimation for heritability, and can provide equivalent estimates of heritability for complex traits after IBD is replaced with IBS (Chen, 2014, 2024).

As we extend the work of Wu and Sankararaman (2018), we similarly assume that

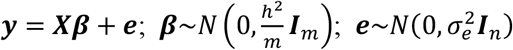

in which ***Y*** is the standardized phenotype of the traits of interest, ***X*** is the standardized genotype matrix of *n* individuals, *m* is the number of double allelic markers, ***β*** is the cumulative effect related to each of the markers, ***e*** is the residual effect, ***I***_*m*_ is an *m* × *m* identity matrix, *h*^2^ is the SNP heritability, and ***I***_*n*_ is an *n* × *n* identity matrix, 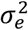 is the residual variance. Under the general assumption for a polygenic trait, it is easy to see that

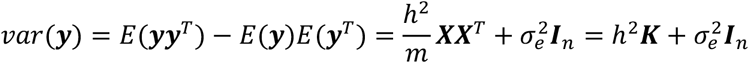

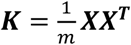 is the genetic relationship matrix (GRM); the moment estimator, or randomized Haseman-Elston regression, is to minimize 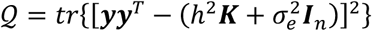. By differentiating *h*^2^ and 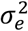 respectively, we have the following normal equations:

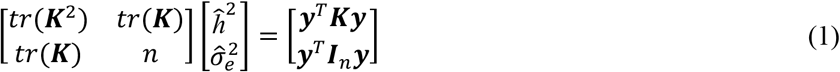

The preliminary estimators for 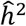 and 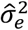 are given as

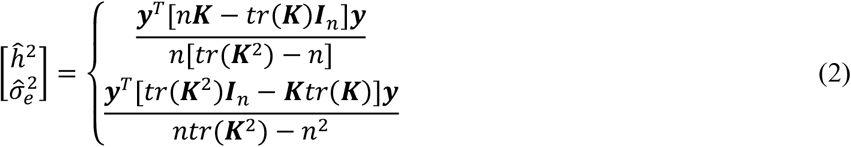

For ease of discussion, we now only focus on the expression without adjustment of covariates. The denominator involves *tr*(***K***^2^), a high-order function for GRM. Alternatively, according to the trace property of a matrix, it can be calculated that 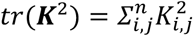, a summation of the square of each element in ***K***. We proved that the expectation of 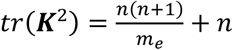, where *m*_*e*_ is the effective number of markers that depicts the average correlations among all genomic markers. Therefore, the expectation for the preliminary estimator of *h*^2^ is 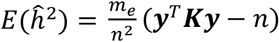. At first glance, it seems inevitable to compute ***K***, the computational cost of which is 𝒪(*n*^2^*m*), a substantial cost given a large sample size, such as for UKB of about 500,000 samples. We obtain the estimate of *tr*(***K***^*c*^) according to the properties of matrix algebra.

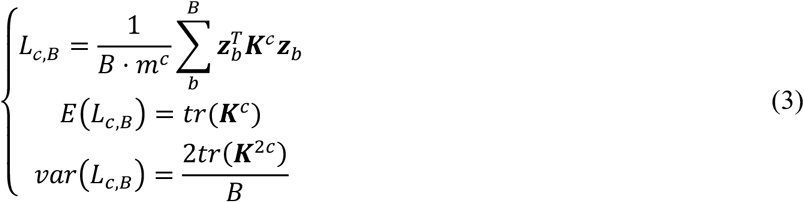

where ***Z***_*b*_ is a vector of length *n* and *B* is the round of iterations. Of note, the sampling variance of of ***L***_2,*B*_is 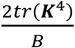. As will be shown below, *tr*(***K***^4^) will be a plugin parameter in the analysis below, and we suggest a robust estimation of 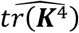 from ***L***_4,*B*_ rather than 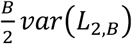. **Eq 3** is the most innovative part of the work of Wu and Sankararaman.

### Randomized estimation for *h*^2^ via RHE-Reg

When there is random mating or little inbreeding, *E*[*tr*(***K***)] = *n*, and substituting the expressions given as **Eq 3** into **Eq 2**, a randomized estimator of heritability is

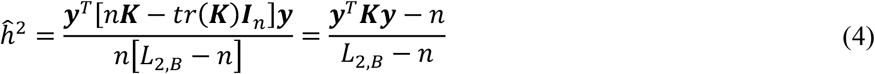

The component 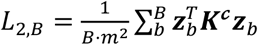 in the denominator is no other than a shuffling nature of the estimation with *B* rounds of resampling.

#### Sampling variance of RHE-reg

The randomized estimator of *h*^2^ can be seen as a ratio of 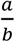 and, according to Delta method, its sampling variance can be expressed as 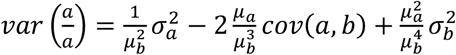, in which the covariance term can be zeroed out in this scenario (Lynch and Walsh, 1998). So, we can obtain

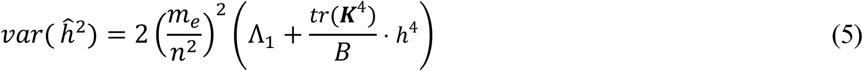

For the definition of Λ_1_ please refer to the section “Estimation for key parameters”. As ***L***_2,*B*_is a random variable, using Tylor approximation *ĥ*^2^ can be obtained by combining **Eq 1**

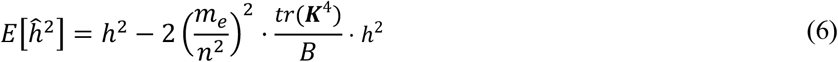

in which the second term is the bias of the RHE-reg estimator. At the same time, we can also find the mean squared error (MSE), the summation of the sampling variance and squared bias, for *ĥ*^2^ as below

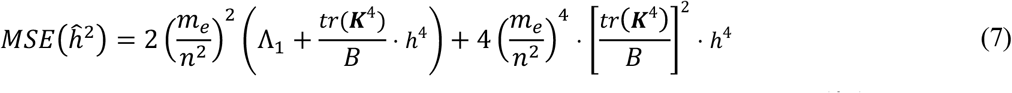

In this polynomial expression, as will be shown in the simulation and real data analysis, *MSE*(*ĥ*^2^) is largely upon the sampling variance and can be further reduced with sufficient iterations (*B*).

#### Constructing test statistics

Given the estimation of heritability, we can construct the *z*-score statistic below:

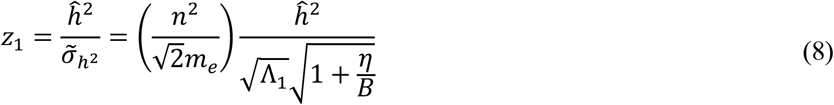

in which 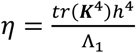, a quantity that will be weighted out with sufficient iterations. Obviously, when *B* is large enough, the optimal *z* score is the following:

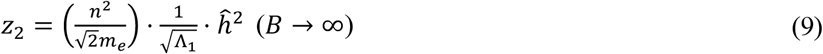

There is an obvious relationship between two z scores in **Eq 8** and **Eq 9**. Given *z*_1_ we can predict optimal test statistic *z*_3_ as below.

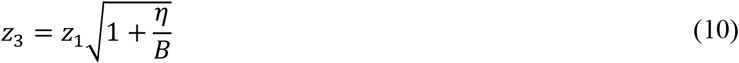

It means that after *B* iteration the expectation of the test statistic is predictable in certain degree.

#### Estimation for key parameters

There are several key quantities/parameters involved in the above equations for RHE-reg, and we present how to estimate them. These parameters are *m*_*e*_ – effective number of markers, *tr*(***K***^4^) the trace of fourth-order GRM, and Λ_1_.

#### Estimation for the effective number of markers (m_e_)

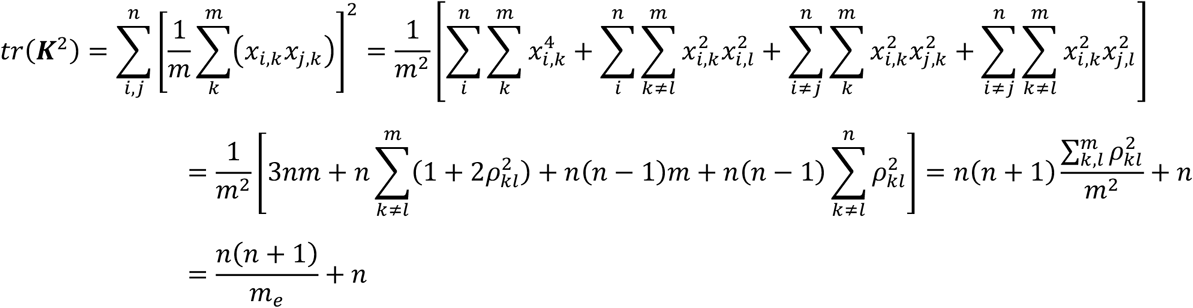

in which 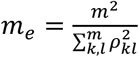 the effective number of markers. Often *m*_*e*_ ≤ *m*, and *m*_*e*_ = *m* if all markers are in linkage equilibrium (see the note of **Table 1**). Here, *m*_*e*_ is a population parameter, a summary statistic that encompasses allelic frequencies and linkage disequilibrium of makers. We consequently propose a randomized algorithm, which estimates *m*_*e*_ as below

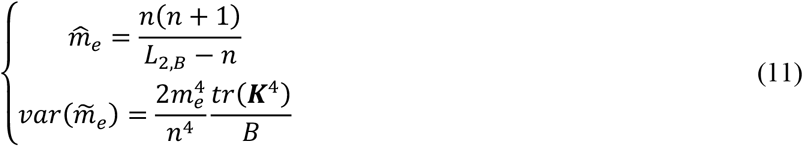

**Table 1:**
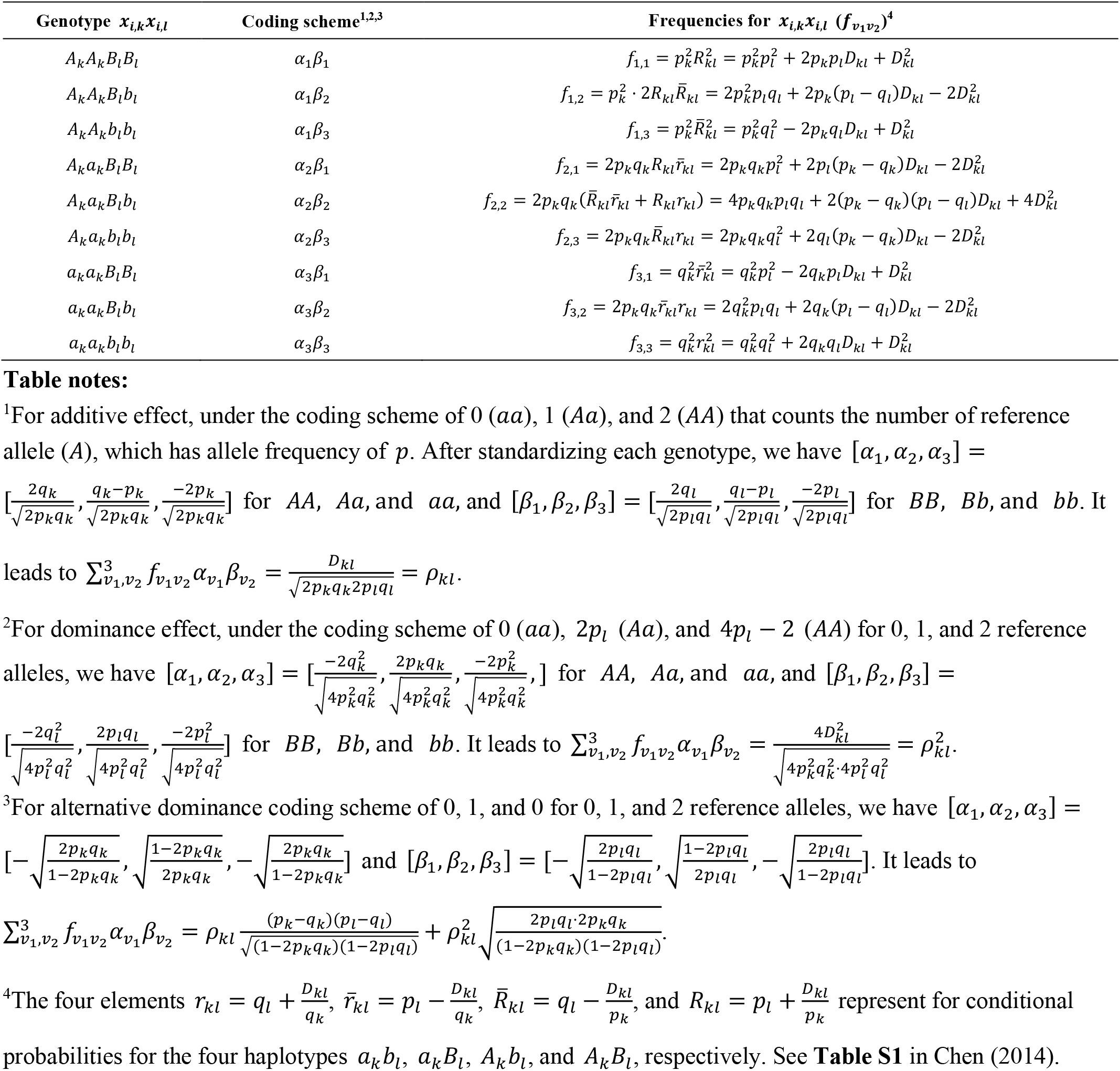
Table for high-order moments for different coding scheme for genotypes.

### Estimation for *tr*(*K*^4^)

The benchmark estimation for *tr*(***K***^4^) is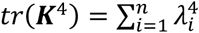, a fourth-order summation of the eigenvalues of ***X***. However, it is computationally expensive when ***X***is large. There are two alternative choices to estimate *tr*(***K***^4^). Method 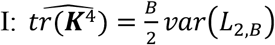, and *B* would affect its precision. Method 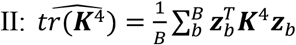, which uses the fourth-order randomized estimation in **Eq 3**. Both Method I and Method II can be realized via **Eq 3**. As will be shown below, Method II provides more stable estimates than Method I.

#### About Λ_1_—high-dimension structure of genetic architecture

For Λ_1_, Λ_1_ = *tr*{[∑(***K*** − ***I***_*n*_)]^2^} = *tr*{∑(***K*** − ***I***_*n*_)∑(***K*** − ***I***_*n*_)}, in which if 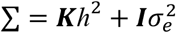 is replaced by ∑ = ***YY***^*T*^, we consequently have

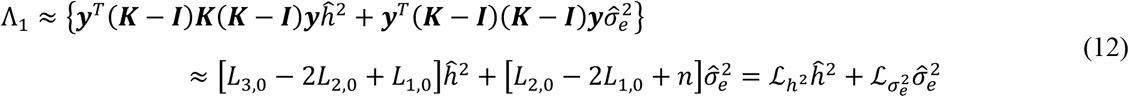

***L***_3,0_ can be estimated as in **Eq 3** if ***Z*** is replaced by ***Y***; it is similarly for ***L***_2,0_ and ***L***_1,0_. They reflect high-dimensional structure between ***Y*** and ***X***. So the sampling variance of *h*^2^ is not only related to *h*^2^ itself, but is eventually upon the high-order structure between ***Y*** and ***X***.

#### About *η* — the term about iteration

We define the ratio for *η* as below

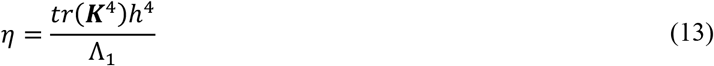

in which *tr*(***K***^4^) can be estimated as 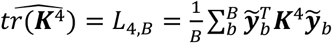 as above. However, it should be noticed that *h*^4^ is a heavy penalty for higher heritability; for example, comparing with *h*^2^ = 0.01, *h*^2^ = 0.1 leads to a 100-fold penalty for the latter in **Eq 13**. Easily, we can estimate *B* if we want to know how many iterations are needed to reach the preset ratio of *η*

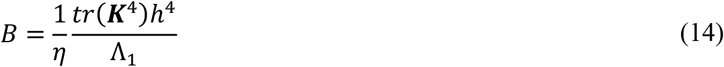

In practice, *η* can take the value of 0.1 or 0.05 as in our simulation and real data analysis below.

#### Extended utilities for distributed datasets

Because datasets are often distributed across institutes, we consequently consider two scenarios for the application of RHE-reg in distributed datasets. As the estimation for *h*^2^ (**Eq 4**) can be split into the numerator and the denominator, the numerator and the denominator are estimated from two different sources. In the other scenario, the whole dataset has been distributed into small slices at different institutes. We call the first scenario the vertical RHE-reg and the latter horizontal RHE-reg.

#### Vertical RHE-reg

Estimation for *h*^2^ can be implemented in summary statistics that the numerator and the denominator can be from different components (Zhou, 2017). The numerator of **Eq 5** can be rewritten as

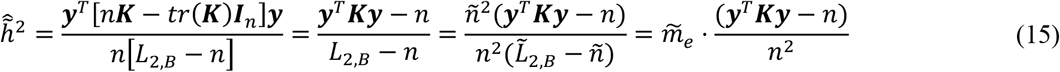

the denominator 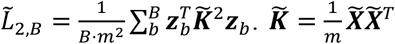, in which 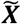 has the dimension of ñ× *m*. So, the sampling variance for 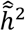 is

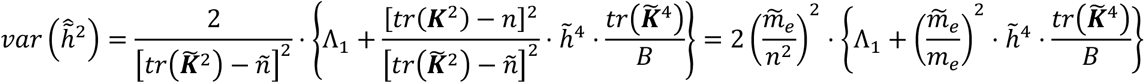

The bias is 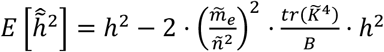, which zeros out when *B* increases. The corresponding test statistic is

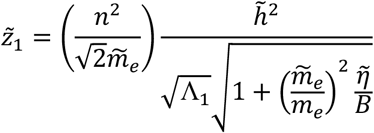

in which 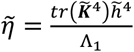. The ratio 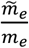, after sufficient iteration, is cancelled out and leads to

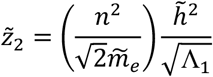

#### Horizontal RHE-reg

For this application, it is assumed that the entire dataset is divided into *c* institutes. Consequently, the whole data ***Y*** and ***X***are distributed in *c* institutes, ***Y***^′^ = [***Y***_1_ ⋮ ***Y***_2_ ⋮ ⋯ ⋮ ***Y***_*c*_] and ***X***^*T*^ = [***X***_**1**_ ⋮ ***X***_2_ ⋮ ⋯ ⋮ ***X***_***c***_]^*T*^. The one only needs to receive the mean and summation of square for each ***Y***_*v*_, and similarly for receiving the allele frequencies of the reference alleles of ***X***_*v*_. So after scaling for ***Y***_*v*_ and ***X***_*v*_,

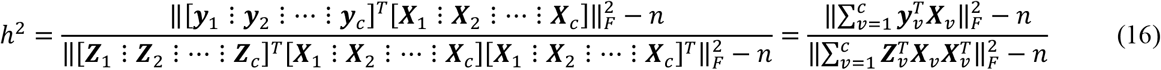

***Z***_*v*_, a *B* × *n*_*v*_ matrix, can be generated from *N*(0,1), by each institute, and consequently independently generate 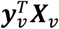 and 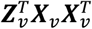 without compromise of privacy. Upon the precision requirement, after *B* rounds of iterations, *η* can be calculated so as to evaluate whether further iterations are needed. Unlike the vertical RHE-reg, the horizontal RHE-reg is identical to the RHE-reg under this simple scenario. An R script is attached for its detailed implementation (**Extended Data 1**).

#### Summary for RHE-reg

Now we discuss some computational issues about RHE-reg. So, eventually *B* will creep into the RHE-reg. The focus here is to investigate how *B* would affect the RHE-reg, in particular the stability of *h*^2^ and z scores. All the above analyses are based on three computational units, ***Y, X***, and *W* – if covariates are taken into account, and the operation between them lead to the whole computational procedure, of which their elementary operations can be implemented hierarchically (**Table 2**). We give an atlas for the computational route. Furthermore, if we have *c* covariates, and the covariate matrix *W* is of *n* × *c* dimensions. After inclusion of the covariates, the equations for stopping rules can be updated accordingly (see **Appendix A**). For fast vector-matrix multiplication, Mailman algorithm is employed here (Liberty and Zucker, 2009). We adapt the implementation of the Mailman algorithm from Agrawal’s fast PCA project (Agrawal *et al*., 2020). It is known that using the Mailman algorithm the vector-matrix multiplication in ***L***_2,*B*_is reduced from 𝒪(*nmB*) to 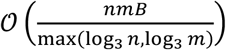. There is no conceptual obstacle to applying the method for genotype data in dosage format, but the Mailman algorithm cannot proceed in such a scenario. We finish the description of the statistical approaches and go to their applications now.

**Table 2.** Analytical results for RHE-reg.

|  | Individual-level data |  | Vertical RHE-reg |  |
| --- | --- | --- | --- | --- |
| $h^2$ estimation | $\begin{cases} h^2 = \frac{\mathbf{y}^T \mathbf{K} \mathbf{y} - n}{L_{2,B} - n} \\ \text{var}(\hat{h}^2) = 2 \left(\frac{m_e}{n^2}\right)^2 \left(\Lambda_1 + \frac{\text{tr}(\mathbf{K}^4)}{B} h^4\right) \end{cases}$ | $\mathbf{z} \begin{cases} z_1 = \left(\frac{n^2}{\sqrt{2}m_e}\right) \frac{\hat{h}^2}{\sqrt{\Lambda_1} \sqrt{1 + \frac{\eta}{B}}} \\ z_2 = \left(\frac{n^2}{\sqrt{2}m_e}\right) \frac{\hat{h}^2}{\sqrt{\Lambda_1}} \end{cases}$ | $\begin{cases} \tilde{h}^2 = \frac{\mathbf{y}^T \mathbf{K} \mathbf{y} - n}{\tilde{L}_{2,B} - \tilde{n}} \\ \text{var}(\tilde{h}^2) = 2 \left(\frac{\tilde{m}_e}{n^2}\right)^2 \left\{\Lambda_1 + \left(\frac{\tilde{m}_e}{m_e}\right)^2 \frac{\text{tr}(\tilde{\mathbf{K}}^4)}{B} h^4\right\} \end{cases}$ | $\mathbf{z} \begin{cases} \tilde{z}_1 = \left(\frac{n^2}{\sqrt{2}\tilde{m}_e}\right) \frac{\tilde{h}^2}{\sqrt{\Lambda_1} \sqrt{1 + \left(\frac{\tilde{m}_e}{m_e}\right)^2 \frac{\tilde{\eta}}{B}}} \\ \tilde{z}_2 = \left(\frac{n^2}{\sqrt{2}\tilde{m}_e}\right) \frac{\tilde{h}^2}{\sqrt{\Lambda_1}} \end{cases}$ |
| Key statistics estimation | $\left\{ \begin{array}{l} \text{Intermediate parameters } \left\{ \Lambda_1 = \mathcal{L}_{h^2} h^2 + \mathcal{L}_{\sigma_e^2} \sigma_e^2, \eta = \frac{\text{tr}(\mathbf{K}^4) \cdot h^4}{\Lambda_1} \right. \\ \text{Randomization} \left\{ \begin{array}{l} \text{tr}(\widehat{\mathbf{K}^2}) = L_{2,B} = \frac{1}{B \cdot m^2} \sum_b \mathbf{z}_b^T \mathbf{K}^2 \mathbf{z}_b \\ \text{tr}(\widehat{\mathbf{K}^4}) = L_{4,B} = \frac{1}{B \cdot m^4} \sum_b \mathbf{z}_b^T \mathbf{K}^4 \mathbf{z}_b \\ \hat{m}_e \left\{ \begin{array}{l} \hat{m}_e = \frac{n(n+1)}{L_{B,2} - n} \\ \text{var}(m_e) = 2 \left(\frac{m_e}{n}\right)^4 \cdot \frac{\text{tr}(\mathbf{K}^4)}{B} \end{array} \right. \end{array} \right. \end{array} \right.$ | | $\left\{ \begin{array}{l} \text{Intermediate parameters } \left\{ \Lambda_1 = \mathcal{L}_{h^2} \tilde{h}^2 + \mathcal{L}_{\sigma_e^2} \tilde{\sigma}_e^2, \tilde{\eta} = \frac{\text{tr}(\tilde{\mathbf{K}}^4) \cdot h^4}{\Lambda_1} \right. \\ \text{Randomization} \left\{ \begin{array}{l} \text{tr}(\widehat{\tilde{\mathbf{K}}^2}) = L_{2,B} = \frac{1}{B \cdot m^2} \sum_b \mathbf{z}_b^T \tilde{\mathbf{K}}^2 \mathbf{z}_b \\ \text{tr}(\widehat{\tilde{\mathbf{K}}^4}) = L_{4,B} = \frac{1}{B \cdot m^4} \sum_b \mathbf{z}_b^T \tilde{\mathbf{K}}^4 \mathbf{z}_b \\ \tilde{m}_e \left\{ \begin{array}{l} \tilde{m}_e = \frac{\tilde{n}(\tilde{n}+1)}{\tilde{L}_{B,2} - \tilde{n}} \\ \text{var}(\tilde{m}_e) = 2 \left(\frac{\tilde{m}_e}{\tilde{n}}\right)^4 \cdot \frac{\text{tr}(\tilde{\mathbf{K}}^4)}{B} \end{array} \right. \end{array} \right. \end{array} \right.$ | |
| | $\hat{\mathcal{L}}_{h^2} = \frac{1}{m^3} \mathbf{y}^T \mathbf{K}^3 \mathbf{y} - \frac{2}{m^2} \mathbf{y}^T \mathbf{K}^2 \mathbf{y} + \frac{1}{m} \mathbf{y}^T \mathbf{K} \mathbf{y}$ $\hat{\mathcal{L}}_{\sigma_e^2} = \frac{1}{m^2} \mathbf{y}^T \mathbf{K}^2 \mathbf{y} - \frac{2}{m} \mathbf{y}^T \mathbf{K} \mathbf{y} + n$ | | | |

Notes: left and right parts of the table give how to implement RHE-reg directly in individual-level data or vertical RHE-reg.

## Results

### Simulation results

We conducted simulations to evaluate the aforementioned theoretical results under various parameters. *h*^2^ was set the value of 0, 0.1, and 0.25, and all SNPs were considered causal after a typical polygenic model; the reference allele frequency was evenly sampled from 0.1∼0.5, and the linkage disequilibrium (Lewontin’s *D* ′) for each pair of consecutive SNPs were *D*′ =0, 0.2, 0.4, 0.6, 0.8 for consecutive SNPs. We set *n* =1,000, 5,000, and 10,000, respectively; *m* =10,000, 50,000, and 100,000. It totally carried out 135 simulation scenarios. For each simulation scenario, we set *B* the value of 10, 20, and 50 in order to find proper *B. n, m* (as well as their allele frequencies), *D*′, and *h*^2^ were considered to investigate how to determine *B*. We investigate and summarize certain properties of RHE-reg in the results below.

### Result 1: Randomized estimation for *tr*(*K*^4^)

As shown in the method section, *tr*(***K***^4^) was appeared as one of the key parameters in determining the performance of the sampling of RHE-reg. The direct estimation of *tr*(***K***^4^) from the eigenvalues of ***K*** was the golden standard, and we consequently compared Method I, 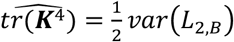, and Method II, 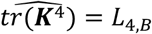, with its direct estimation. As shown in **Figure 1**, the above 135 simulation scenarios were compared with the direct estimation for 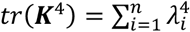. For Method I, increasing *B* from 10 to 50 could increase the precision of the estimation. In contrast, Method II showed very consistent and high precision for the estimation of *tr*(***K***^4^) regardless of the sample size, an increasing of *B* from 10 to 50 did not help improve precision. The advantage of Method II was probably because ***L***_4,*B*_estimated *tr*(***K***^4^) as its mean, whereas *var*(***L***_2,*y*_) as its sampling variance. So, hereafter we used Method II ***L***_4,*B*_to estimate *tr*(***K***^4^).

**Figure 1.**
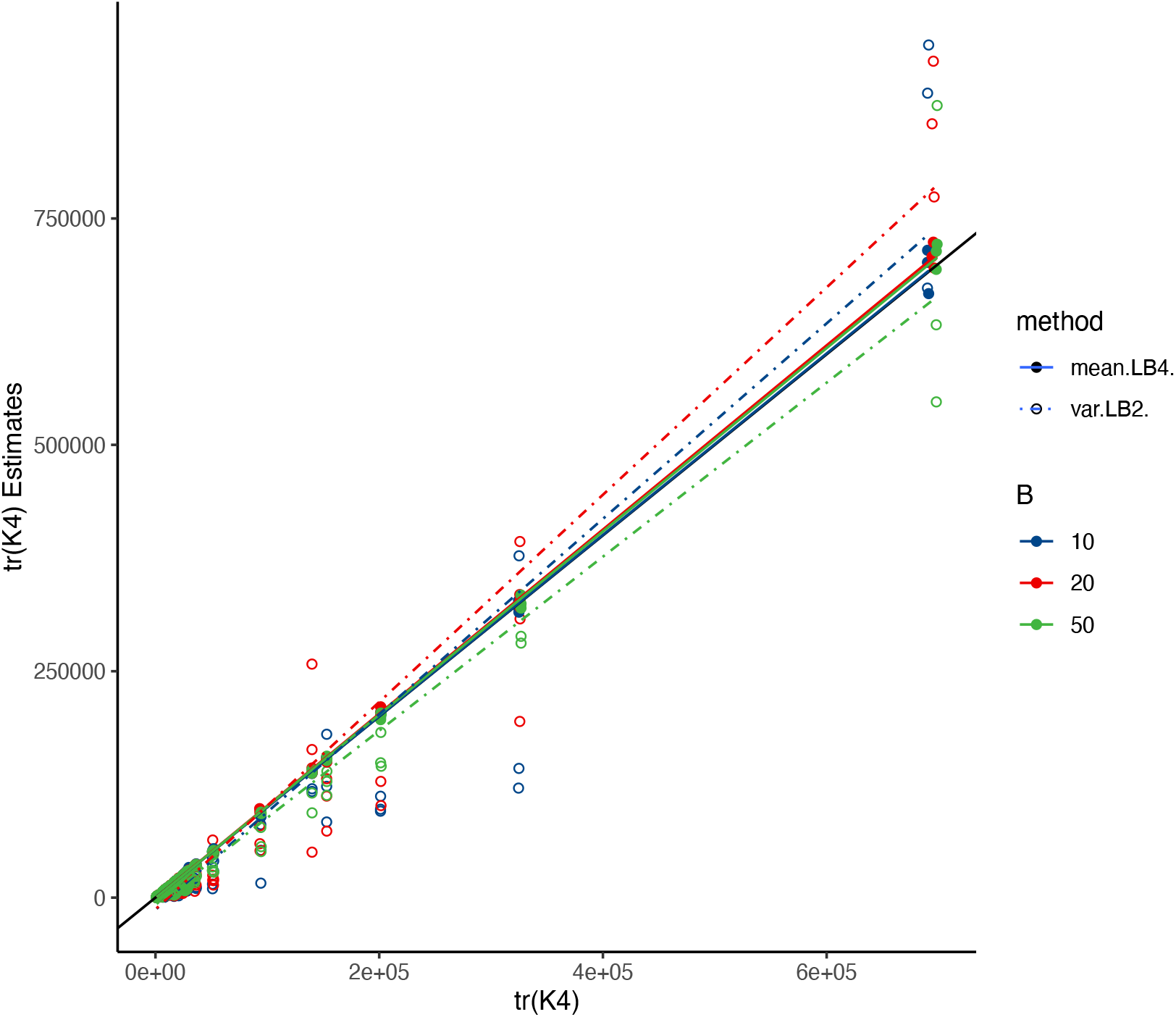
Comparison for the estimation of *tr*(***K***^4^). The x-axis represents benchmark estimation for *tr*(***K***^4^) directly, and y-axis represents the estimation of *tr*(***K***^4^) using Method I or Method II respectively. The diagonal line is for comparison. Each fitted line shows the correlation between all 135 estimations with their benchmark estimation *tr*(***K***^4^).

### Result 2: MSE of RHE-reg

For ease of presentation, we reorganize **Eq 7** by denoting with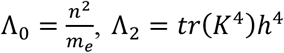, and Λ_3_ = *tr*(***K***^4^), which gives 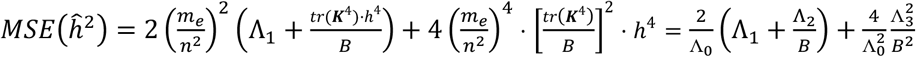. We compared these three components 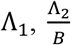, and 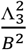, respectively. We only illustrated the result for *n* = 5,000 and 10,000, respectively, because *n* = 1,000 was too small a sample size here.

As Λ_1_ depended little on *B*, we could use Λ_(_ as a benchmark to examine how *B* could shrink MSE by reducing 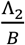 and 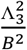. The regression coefficient fitted in each plot in the first row of **Figure 2** reflected how quickly *B* could reduce 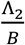, and the regression coefficient also reflected the quantity of *η* after *B* iterations. A much decreasing slope of the fitted regression indicated much less influence of Λ_2_ as well as Λ_3_ to MSE. In contrast, Λ_3_ was seemed a less important term in the MSE of *h*^2^ via RHE-reg, and vanished much faster than Λ_2_. Furthermore, a much higher *h*^2^ took a much greater *B* in comparing the regressions across all scenarios. LD did not play an important role in determining MSE.

**Figure 2.**
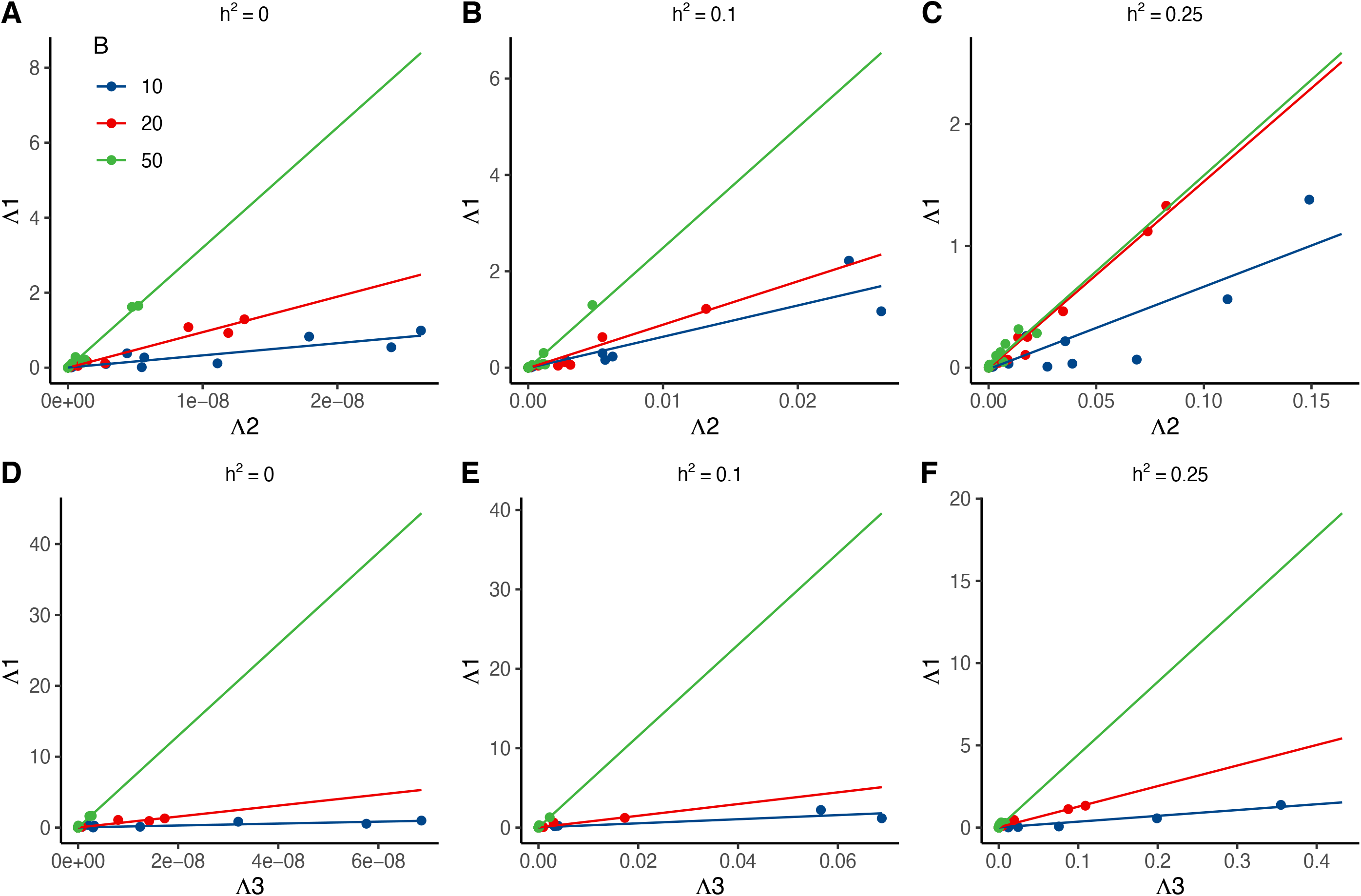
Evaluation for the MSE of RHE-reg under the different simulation scenarios. The first row represents the comparison between Λ_2_ and Λ_1_ under different *h*^2^ and *B*, and the second row represents the comparison between Λ_3_ and Λ_1_. The flatter the fitted line is, the smaller Λ_2_ or Λ_3_ is compared with Λ_1_

### Result 3: Randomized estimation for *h*^2^ and *z*-score

In result 3, we studied how *B* could influence *h*^2^ and its *z*-score. As the sampling variance of *h*^2^ was reciprocal to the sample size 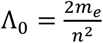 under the null hypothesis and *B*, it was obviously to see in simulation that: greater *n*, and greater *B* would help to bring out a more stable estimation for *h*^2^ (**Figure 3A-C**). If we employed *ĥ*^2^ from *B* = 50 as the benchmark, when sample size *n* = 10,000, there was very high consistent estimation for *ĥ*^2^ even *B* = 10 and *B* = 20 (**Figure 3C**).

**Figure 3.**
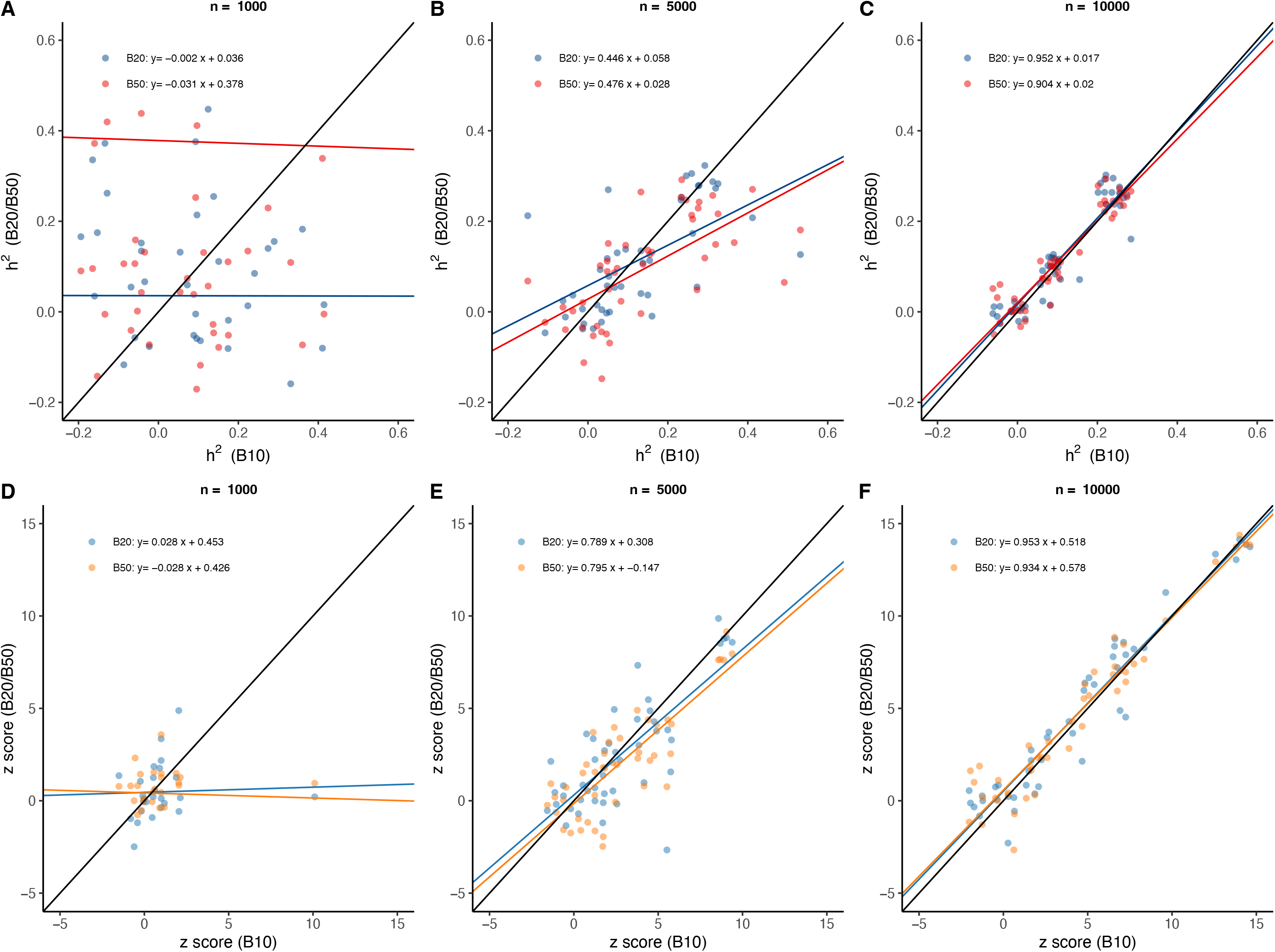
Estimation of *h*^2^ and z-score after different *B*. Each plot illustrates the comparison of the estimated heritability **(A-C**) and *z*-score (**D-F**) given *B* = 10 (x-axis) vs *B* =20 and 50 (y-axis) under different sample size *n* = 1,000, 5,000, and 10,000, respectively. The black solid line is the reference line of y=x, and the coloured solid line is the fitted regression, which is printed in each plot.

The availability of the *z* score of the estimated heritability was important for statistical inference. We evaluated the influence of *B* in determining the performance of the randomized algorithm (**Figure 3D-F**). It was known from the above analysis 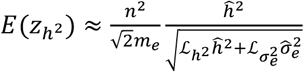, so when the estimation of *h*^2^ became stable the test statistic was stable too. So, *z*_1_ was relative stable when *n* = 5,000 (**Figure 3E**) or *n* = 10,000 (**Figure 3F**). When the sample size was sufficiently large, a few iteration could guarantee high accuracy of the estimation.

### Result 4: Application of horizontal RHE-reg

This study was to estimate heritability for distributed data as exact as a single piece of data. Two cohorts with *n*_1_= 4,000 and *n*_2_ = 6,000 individuals, respectively, were generated to verify h-RHE-reg. *h*^2^ was set the value of 0, 0.1, and 0.25, respectively. The effects of all *m* = 10,000 SNPs were sampled from the distribution 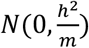. Heritability and z scores were estimated using individual-level RHE-reg as well as h-RHE-reg. *B* was set of 10, 20, and 50. The genotypes of the two simulated cohorts were standardized by 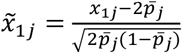 and 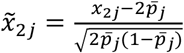 for the *j*-th locus, where 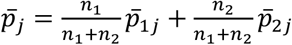 was the average allele frequency. The phenotypes of the two cohorts were standardized by 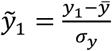 and 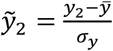, where 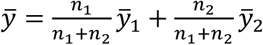 and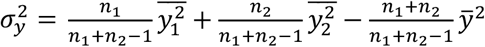. Each simulation scenario had 10 repeats.

The estimates of heritability and its z score were consistent using individual-level RHE-reg and h-RHE-reg in all scenarios when the random vectors were the same (**Figure 4**).

**Figure 4.**
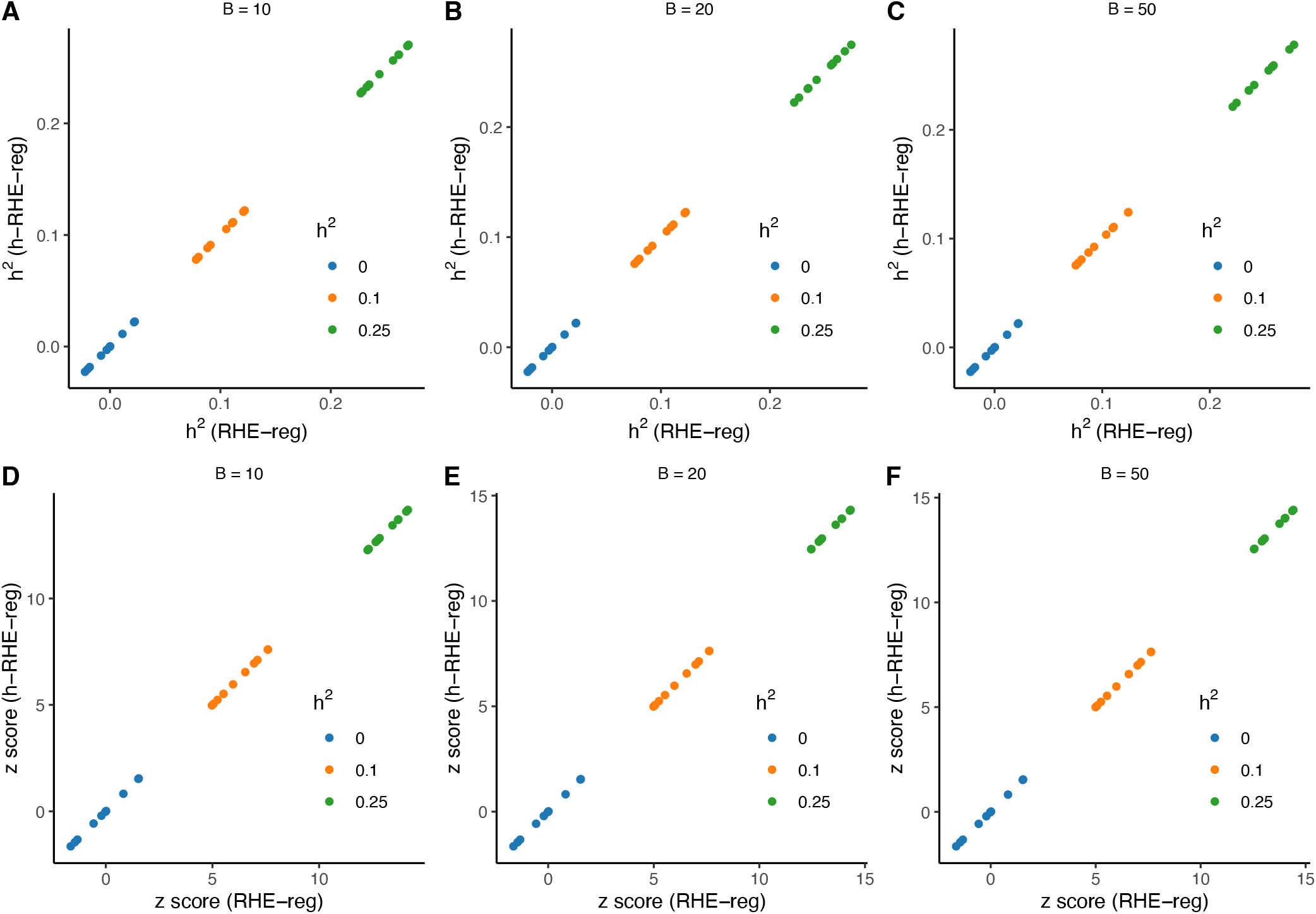
Application of horizontal RHE-reg in simulation studies. Estimated heritability (**A-C**) and z-scores (**D-F**) through RHE-reg (x axis) and h-RHE-reg (y axis) are given under different *B* =10, 20, and 50.

### Real data analysis for UK Biobank

We chose the unrelated 292,223 British white who have no kinship found, as indicated by the genetic kinship provided in the UK Biobank (field 22021) for real data test (Bycroft *et al*., 2018). After quality control, the inclusion criteria were: MAF > 0.01, missing call rate < 0.05 and Hardy-Weinberg proportion test *p*-value > 1e-6, whose genotype call rate > 0.95, and 525,460 autosome SNPs with were included for analysis. We estimated heritability of the 81 quantitative traits, and included the top two principal components and sex as covariates.

We used two strategies to estimate heritability. In strategy I, denoted as *B+* strategy hereafter, we set *B*_0_ = 10 as a warm-up step to evaluate *tr*(***K***^4^) and *η* was set of 0.05. After the warm-up of *B*_0_ iteration, we then increased iteration by a step of 10, We then estimate final realized *η, m*_*e*_, *h*^2^, and three kinds of z scores until the convergence ratio of 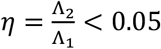 (Λ_2_ was as defined in **Result 2**); however, we set a hard stop for *B*_1_ = 200 even if *η* was still greater than 0.05. In strategy II, we directly set *B*_0_ = 10, 20, or 50 without further considering additional iteration anymore, and consequently denoted as *B*10, *B*20, and *B*50 strategies hereafter.

### Comparison of the UKB results between strategy I and II

Even a couple of traits were set to take a hard stop because their *B*_1_ were greater than 200, the estimated 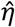 for the 81 traits was 0.0518, which was very close to the preset *η* = 0.05 (**Figure 5 A1**). It indicated that our theory worked to control the precision of the sampling variance of RHE-reg. The trait “age of diabetes diagnosed” had *h*^2^ = 1.21 ± 0.533, extremely large standard error compared to other traits, because of its smallest sample size of *n* = 12,658 (**Figure 5B1**). In **Figure 5 C1**, we got three *z* scores, which are 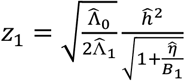 score directly calculated given *B*_1_ iterations (green colored, via **Eq 8**), the optimal z score 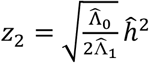 when *B* was infinitive (blue colored, via **Eq 9**), and the predicted 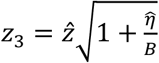 score (pink colored, via **Eq 10**).

**Figure 5.**
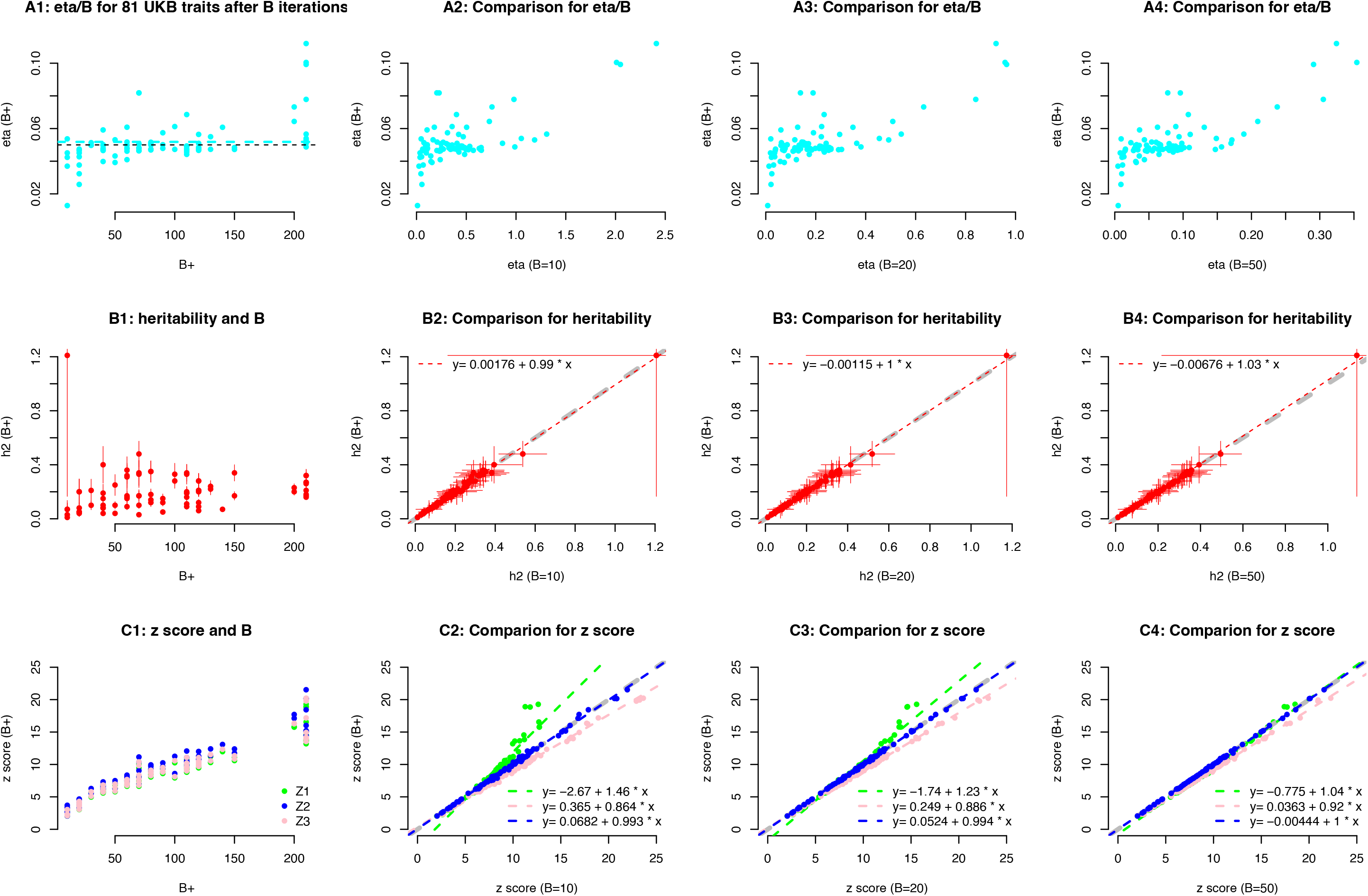
Randomized estimation for heritability for UK Biobank 81 quantitative traits. A1-D1) The performance of RHE-reg given the respective *B* number for each trait, η, effective number of markers using randomized estimation (*m*_*e*_), *ĥ*^2^ (the vertical line covers 95% confidence interval), and *z* scores estimated in three methods. Three z scores are plotted, the green colored z scores are directly estimated given *B* iterations for each trait (Eq 8), the pink colored z scores are optimal z score (Eq 9), and the blue colored z scores are directly estimated given 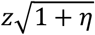(Eq 10). A2-A4) Comparison for η between that of *B* + and *B* = 10, 20, and 30, respectively. B2-B4) Comparison for *m*_*e*_ between that of *B* + and *B* = 10, 20, and 30, respectively; the vertical and horizontal lines are the means of *m*_*e*_ from x-axis and y-axis, respectively. C2-C4) Comparison for *h*^2^ between *B* + and *B* = 10, 20, and 30, respectively; the fitted lines is printed on the top left corner of each plot. D2-D4) Comparison for the three pairwise *z* scores. The green colored z scores are estimated in **Eq 8** given B+ and the number of B as shown on the x-axis label, the pink colored z score are estimated in **Eq 9**, and blue colored ones in **Eq 10**, respectively.

For comparison, we examined the corresponding statistics that were estimated under *B*10, *B*20, and *B*50, respectively. In strategy II, A larger *B*_0_ led a smaller *η* as expected (**Figure 5 A2-4**). Interestingly, regardless the change of *B*_0_ in strategy II, 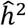 were very consistent to those estimated from strategy I, as shown that the fitted regression lines were very close to 1 (**Figure 5 B2-B4**). Three types of *z* scores were compared (**Figure 5 C2-C4**), and the optimal *z* scores from both strategies were nearly perfect (blue points and blue dashed lines). Then, as shown in **Figure 5 C4**, the three kinds of *z* scores were nearly completely matched.

In addition, the estimates were also consistent with our previous results using a less efficient method (Xu *et al*., 2021), and see **Extended Data 2** for more details. The heritability estimated by the randomization algorithm exhibited a high degree of correlation (Pearson’s correlation coefficient 0.77) with the previous estimates for 81 traits. Compared to the previous results, the *m*_*e*_ was nearly consistent with GRM-based estimates, and is with averaged 1.38% deviation after 10 iterations and further decreased to 1.23% deviation after 50 iterations (**Extended Data 2**).

### Application of vertical RHE-reg

We split each UKB trait evenly into halves to test the v-RHE-reg. Of each trait, its heritability and *z* score tests could be constructed within each split and between each split by exchanging the ***L***_*B*_ estimation, and consequently brought out v-RHE-reg. As shown in **Eq 15**, we compared the result for *B*=10, 20, and 50, respectively, and observed consistent results between split 1 and split 2, and between split 1/2 and split 2/1. So, we had four estimators as below

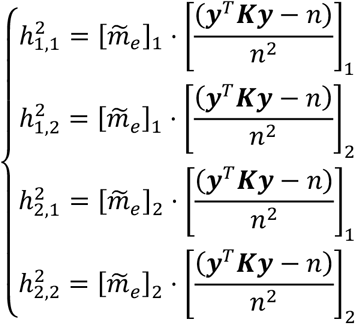

**Figure 6** showed the results of these four estimators under different *B*. It illustrated that pairwise estimates 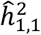 against 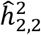, and 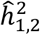 against 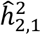, and as observed the pairwise estimates were quite consistent with each other both within and between splits.

**Figure 6.**
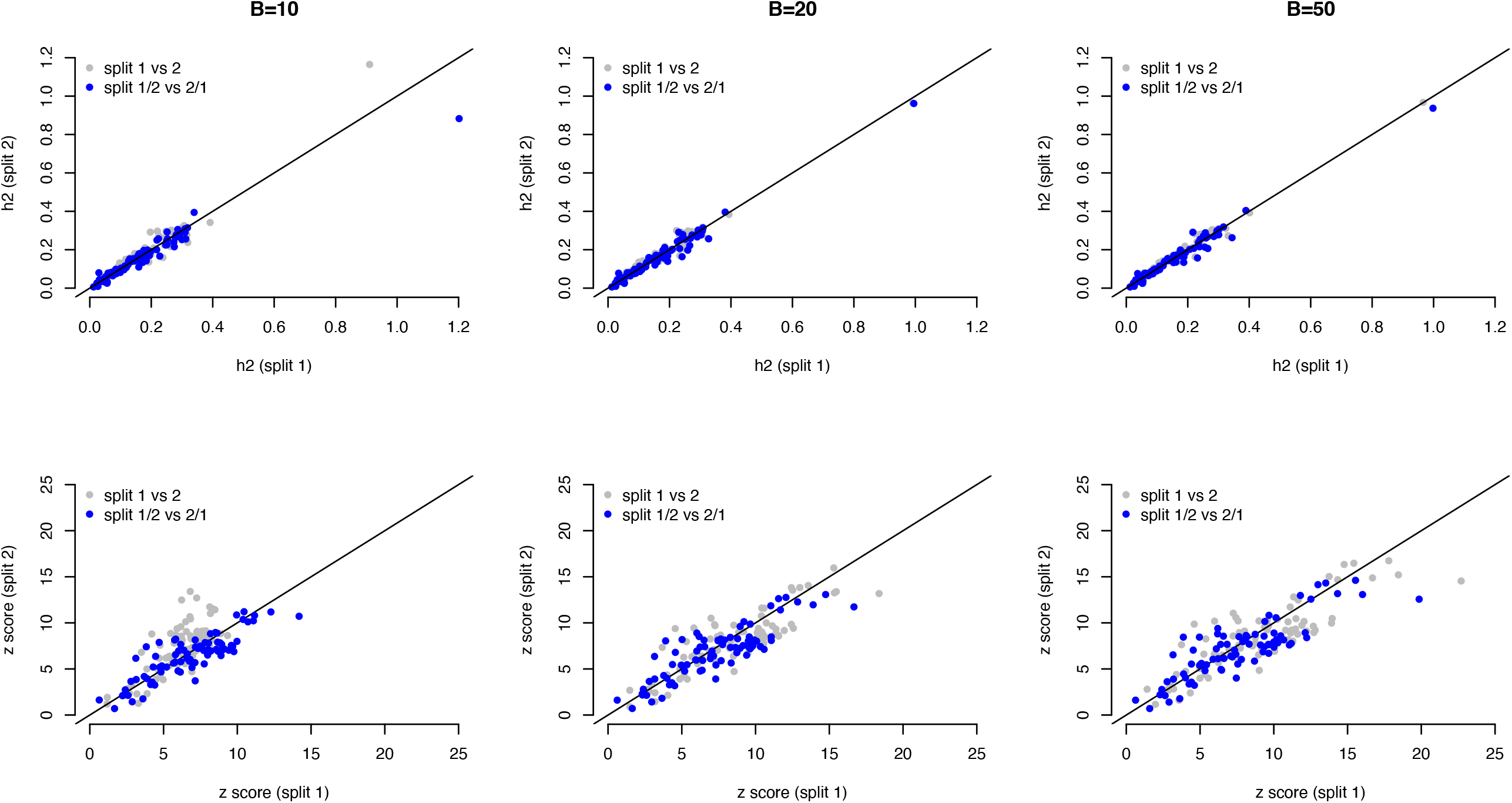
Application of v-RHE-reg for 81 quantitative traits in UKB. The top and the bottom row represent heritability and z score respectively. Each column illustrates results after *B* iterations. In each plot, the coordinate for a grey point is heritability/z-score estimated from split 1 (x-axis) and split 2 (y-axis).

## Discussion

The presented study is developed on the randomized Haseman-Elston regression for the estimation of SNP-heritability proposed recently by Wu and Sankararaman (2018). They very smartly used a randomization approach, which significantly reduces the computational cost in estimating *tr*(***K***^2^) from 𝒪(*n*^2^*m*) to 𝒪(*nmB*). However, the drawbacks of their method may be its unclear property for *B*, which further leads to obscure sampling variance of the estimated heritability. As discussed in a recent review, it has been obscure in the original RHE-reg since no closed-form solutions were provided to quantify the connection between *B* and the estimation procedure (Tang *et al*., 2022). After integrating analytical results for Haseman-Elston regression into this randomized framework (Chen, 2014), we present here a close-form solution for RHE-reg. Having provided the sampling variance, we are able to evaluate how *B* influences the estimation procedure of RHE-reg precisely. In particular, a key element that is related to the sampling variance of ***L***_2,*B*_, which is proportional to 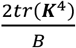. It should be noticed 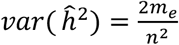 under the null hypothesis that *h*^2^ = 0. The quantity of 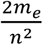 is identical to the sampling variance of REML under the null hypothesis or that of modified Haseman-Elston regression (Visscher *et al*., 2014; Chen, 2014; Zhou, 2017).

It is straightforward to apply the estimation procedure for the estimation of dominance variance components both for individual-level data and summary statistics. The only update of the equation 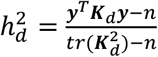 is to replace ***K*** with 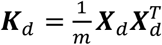. For each SNP, ***x***_*i,d*_ is coded 0, 2p, and 4p-2 for the genotype that counts 0, 1 and 2 reference alleles; and furthermore, ***X***_*d*_ is further scale by 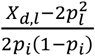 (Zhu *et al*., 2015; Vitezica *et al*., 2017). So for a pair of individual *i* and *j*, 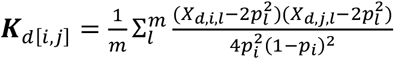. After replacing ***K*** with ***K***_*d*_, all the above estimation procedure can be applied for 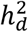. Furthermore,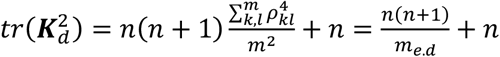. The effective number of markers in terms of ***X***_*d*_ is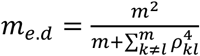, a tetradric form of LD for a pair of SNPs.

A nature extension of the method is to include multi-component, such as for the estimation for each chromosome. It is obvious that the method for deriving sampling variance should be extended for multi-components estimation if their corresponding ***X***_*i*_ and ***X***_*j*_ are in global linkage, or nearly, equilibrium, which is often the case for human populations (Huang *et al*., 2023). Much advanced numerical tools, such as condition numbers, are needed to evaluate the approximation of the randomized algorithm (Horn and Johnson, 1994).

In summary, the purpose of the present study is two-fold. First, we provide a method to balance iteration and precision of estimation, and an improved implementation of RHE-reg is realized. Secondly, we extend RHE-reg into the estimation of SNP-heritability for distributed data, which uses the controlled *B* to synchronize the estimation across datasets. With increasing genomic cohorts but distributed in different institutes, it is now a trend to propose computational solutions without compromising privacy (Elhussein *et al*., 2024). The enhanced RHE-reg framework can consequently have computational and analytical merits, and, as demonstrated, we further extend its utilities such as vertical- and horizontal RHE-reg, as demonstrated in this study. Given the increasing cry for genomic privacy, both vertical and horizontal RHE-reg will be meaningful in securing genomic information. However, given its traditionally very quantitative origin of statistical genetics, statistical routines may have competing, if not superior, solutions than those derived from available information technology (Wang *et al*., 2022; Zhang *et al*., 2024).

## Supporting information

ExtendData1-R code demo

ExtendData2

## Acknowledgements

We thank the participants of the included cohorts and of UK Biobank for making this work possible (UKB application 41376). This work was supported by the National Natural Science Foundation of China (31771392 to GBC, and 32102503 to ZZ), Shenzhen Basic Research Foundation (20220818100717002 to SL), Guangdong Basic and Applied Basic Research Foundation (2022B1515120080 to SL). The funders had no role in study design, data collection and analysis, decision to publish, or preparation of the manuscript.

## Appendix A

### When there is adjustment for covariates *W*

We can derive the equation below with the inclusion of covariates.

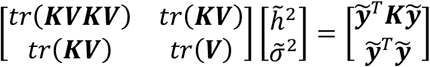

In which ***V*** = ***I*** − *W*(*W*^*T*^*W*)^−1(^*W*^*T*^ and the covariance matrix *W* is *n* × *c* matrix, MSE becomes

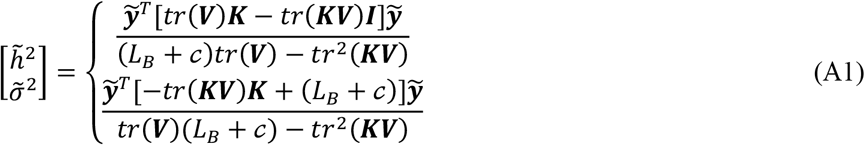

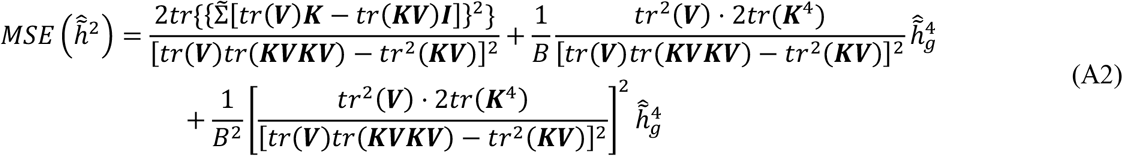

in which ***V*** = ***I***_*n*_ − *W*(*W*^*T*^*W*)^−1(^*W*^*T*^. And consequently,

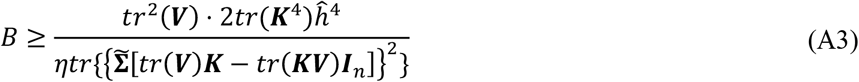

When there is covariance matrix *W*, which is *n* × *c* matrix, MSE becomes

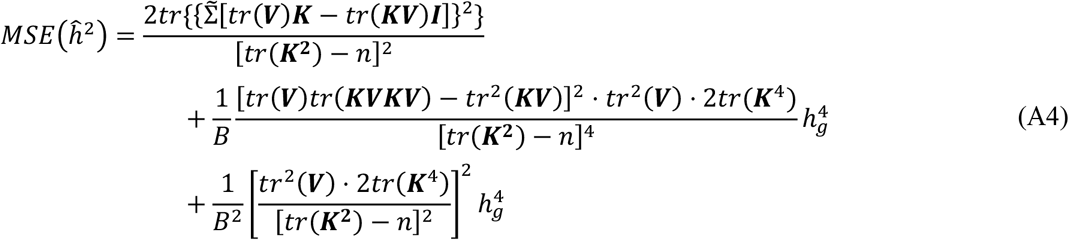

in which ***V*** = ***I***_*n*_ − *W*(*W*^*T*^*W*)^−1(^*W*^*T*^. And consequently,

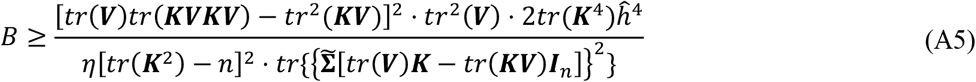

And a hybrid one

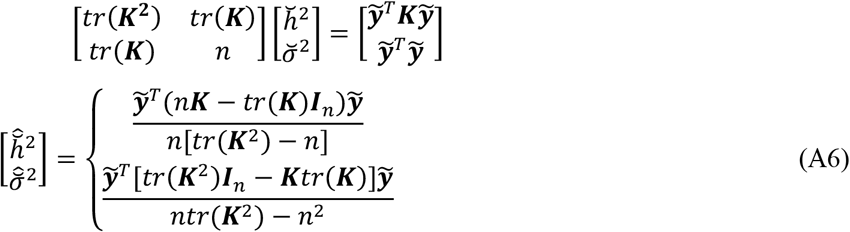

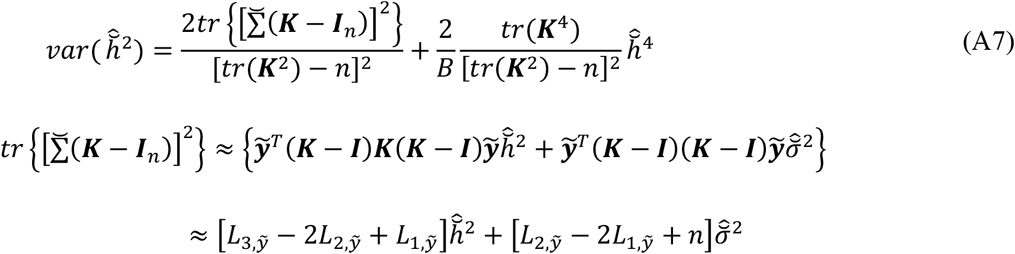

At the same time, we can also find the mean squared error (*MSE*) for *ĥ*^2^ as below

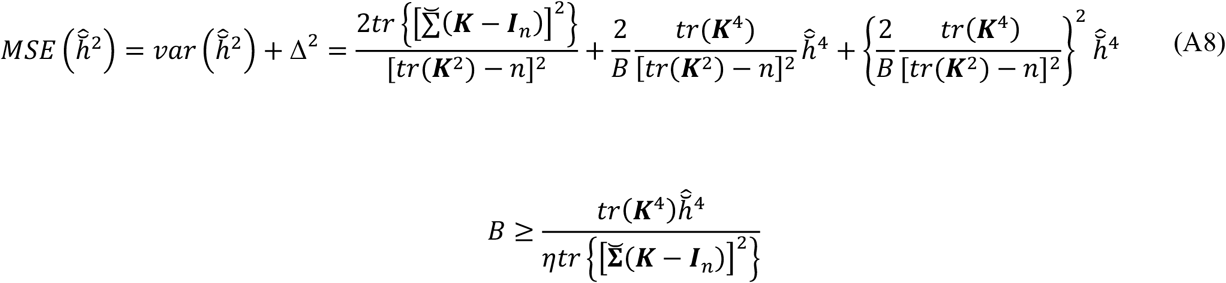

