## Supplementary material for "Analytical and computational solution for the estimation of SNP-heritability in biobank-scale and distributed datasets": ExtendData1-R code demo: simulation.R

```
source("source.R")
library(dplyr)
set.seed(421)
rep = 10

## Settings ----
t0 = Sys.time()
args = commandArgs(T)
B = as.numeric(args[1])  # B = 10
h2 = as.numeric(args[2])  # h2 = 0.1
m = 10000
n1 = 4000
n2 = 6000
n = n1+n2

## Data generation ----
freq = runif(m, 0.1, 0.5)
# Simulate genotype
X1 = GenerateGeno(freq, n1)
X2 = GenerateGeno(freq, n2)

rst = list()
for(r in 1:rep){
  seed = 421 + r
  set.seed(seed)

  # Simulate phenotype
  b = rnorm(m, sd=sqrt(h2/m))
  g1 = X1 %*% b
  g2 = X2 %*% b
  sigma_e = ifelse(h2==0, 1, (1/h2-1)*var(c(g1,g2)))
  y1 = g1 + rnorm(n1, sd=sqrt(sigma_e))
  y2 = g2 + rnorm(n2, sd=sqrt(sigma_e))
  
  ## h-RHE-reg ----
  ### Data standardization ----
  # Genotype Standardization
  w1 = n1/(n1+n2)
  w2 = n2/(n1+n2)
  p1_bar = colMeans(X1)/2 # summary
  p2_bar = colMeans(X2)/2 # summary
  p_bar = w1*p1_bar + w2*p2_bar
  X1s = t(apply(X1, 1, function(x) {(x-2*p_bar)/sqrt(2*p_bar*(1-p_bar))}))
  X2s = t(apply(X2, 1, function(x) {(x-2*p_bar)/sqrt(2*p_bar*(1-p_bar))}))
  # Phenotype standardization
  y_bar = w1*mean(y1) + w2*mean(y2)
  y_var = n1/(n1+n2-1)*mean(y1^2) + n2/(n1+n2-1)*mean(y2^2) - (n1+n2)/(n1+n2-1)*y_bar^2
  y1s = (y1-y_bar)/sqrt(y_var) # approx n = n - 1
  y2s = (y2-y_bar)/sqrt(y_var)

  ### Estimation ----
  ### Denominator - Lb
  seed = 421 + r
  set.seed(seed)
  Lb2 = array(0, dim = B)
  Lb4 = array(0, dim = B)
  for(i in 1:B){
    z = rnorm(n)
    z1 = z[1:n1]
    z2 = z[(n1+1):n]
    zx1 = z1 %*% X1s
    zx2 = z2 %*% X2s
    zx_sum = zx1 + zx2
    zxx1 = zx_sum %*% t(X1s)
    zxx2 = zx_sum %*% t(X2s)
    Lb2[i] = sum(c(zxx1, zxx2)^2)/m^2
    
    zxxx1 = zxx1 %*% X1s
    zxxx2 = zxx2 %*% X2s
    zxxx_sum = zxxx1 + zxxx2
    zxxxx1 = zxxx_sum %*% t(X1s)
    zxxxx2 = zxxx_sum %*% t(X2s)
    Lb4[i] = sum(c(zxxxx1, zxxxx2)^2)/m^4
  }
  me = n*(n+1)/(mean(Lb2)-n)
  
  ### Numerator - Ly1
  yx1 = t(y1s) %*% X1s
  yx2 = t(y2s) %*% X2s
  yx_sum = yx1 + yx2
  Ly1 = sum(yx_sum^2)/m
  
  yxx1 = yx_sum %*% t(X1s)
  yxx2 = yx_sum %*% t(X2s)
  Ly2 = sum(c(yxx1, yxx2)^2)/m^2
  
  yxxx1 = yxx1 %*% X1s
  yxxx2 = yxx2 %*% X2s
  yxxx_sum = yxxx1 + yxxx2
  Ly3 = sum(yxxx_sum^2)/m^3
  
  ## Heritability estimation
  h2hat = (Ly1-n)/(mean(Lb2)-n)
  print(h2hat)
  
  ## Mediators estimation
  lambda1 = (Ly3-2*Ly2+Ly1)*h2hat + (Ly2-2*Ly1+n)*(1-h2hat)
  eta = mean(Lb4)*h2hat^2/lambda1
  
  ## Statistics
  z = (n^2)*h2hat/sqrt(2*lambda1*(1+eta/B))/me
  var = 2*(me/n^2)^2*(lambda1 + mean(Lb4)*h2hat^4/B)
  bias = 2*(me/n^2)^2*(mean(Lb4)*h2hat^2/B)
  
  rst[[2*r-1]] = data.frame(method = "h-RHE-reg", rep = r, h2 = h2, h2hat = h2hat, var = var, z = z, bias = bias, B = B, 
                        me = me, trK4 = mean(Lb4), lambda1 = lambda1, eta = eta)

  ## v-RHE-reg ----
  ### Data standardization ----
  X = rbind(X1, X2)
  y = c(y1,y2)
  p = colMeans(X)/2
  Xs = t(apply(X, 1, function(x) {(x-2*p)/sqrt(2*p*(1-p))}))
  ys = as.matrix((y-mean(y))/sqrt(var(y)))

  ### Estimation ----
  ### Denominator - Lb
  Lb2 = randomization_Lb2(Xs, B, seed = seed)
  Lb4 = randomization_Lb4(Xs, B, seed = seed)
  me = n*(n+1)/(mean(Lb2)-n)
  
  ### Numerator - Ly1
  Ly1 = randomization_Ly1(Xs, ys)
  Ly2 = randomization_Ly2(Xs, ys)
  Ly3 = randomization_Ly3(Xs, ys)
  
  ## Heritability estimation
  h2hat = (Ly1-n)/(mean(Lb2)-n)
  print(h2hat)
  
  ## Mediators estimation
  lambda1 = (Ly3-2*Ly2+Ly1)*h2hat + (Ly2-2*Ly1+n)*(1-h2hat)
  eta = mean(Lb4)*h2hat^2/lambda1
  
  ## Statistics
  z = (n^2)*h2hat/sqrt(2*lambda1*(1+eta/B))/me
  var = 2*(me/n^2)^2*(lambda1 + mean(Lb4)*h2hat^4/B)
  bias = 2*(me/n^2)^2*(mean(Lb4)*h2hat^2/B)
  
  rst[[2*r]] = data.frame(method = "RHE-reg", rep = r, h2 = h2, h2hat = h2hat, var = var, z = z, bias = bias, B = B, 
                        me = me, trK4 = mean(Lb4), lambda1 = lambda1, eta = eta)
}

df = bind_rows(rst)

save(df, file = paste0("simulation-h2-",h2,"-B-",B,".Rdata"))


t1 = Sys.time()
print(t1-t0)

## Plot ----
# library(ggplot2)
# library(ggsci)
# library(egg)
# df0 = data.frame()
# for(h2 in c(0, 0.1, 0.25)){
#   for(B in c(10,20,50)){
#       load(paste0("simulation-h2-",h2,"-B-",B,".Rdata"))
#       df0 = rbind(df0, df)
#   }
# }
# 
# a = 0
# p = list()
# q = list()
# for(b in c(10,20,50)){
#   a = a + 1
#   df = df0 %>% filter(B==b) %>% select(method, rep, h2, h2hat) %>%
#     group_by(rep, h2) %>%
#     spread(key = method, value = h2hat)
#   df$rep = as.factor(df$rep)
#   df$h2 = as.factor(df$h2)
#   p[[a]] = ggplot(df, aes(x = `RHE-reg`, y = `h-RHE-reg`, color = h2)) +
#     geom_point() +
#     labs(title = paste0("B = ",b),
#          x = bquote(h^2~"(RHE-reg)"), y = bquote(h^2~"(h-RHE-reg)")) +
#     scale_color_d3(name = bquote(h^2)) +
#     theme_classic() +
#     theme(legend.position = c(0.8,0.2),
#           plot.title = element_text(size = 9, hjust = 0.5))
# 
#   df = df0 %>% filter(B==b) %>% select(method, rep, h2, z) %>%
#     group_by(rep, h2) %>%
#     spread(key = method, value = z)
#   df$rep = as.factor(df$rep)
#   df$h2 = as.factor(df$h2)
#   q[[a]] = ggplot(df, aes(x = `RHE-reg`, y = `h-RHE-reg`, color = h2)) +
#     geom_point() +
#     labs(title = paste0("B = ",b), x = "z score (RHE-reg)", y = "z score (h-RHE-reg)") +
#     scale_color_d3(name = bquote(h^2)) +
#     theme_classic() +
#     theme(legend.position = c(0.8,0.2),
#           plot.title = element_text(size = 9, hjust = 0.5))
# }
# 
# pdf("simulation-n10000.pdf", width = 10, height = 7, onefile = F)
# ggarrange(p[[1]],p[[2]],p[[3]],
#           q[[1]],q[[2]],q[[3]],nrow = 2, ncol = 3, labels = c("A","B","C","D","E","F"))
# dev.off()
```
