## Supplementary material for "Analytical and computational solution for the estimation of SNP-heritability in biobank-scale and distributed datasets": ExtendData1-R code demo: source.R

```
randomization_Lb2 <- function(X, B, seed){
  # X: scaled genotype
  # B: iteration time
  set.seed(seed)
  
  n = nrow(X)
  m = ncol(X)
  
  Lb2=array(0, dim=B)
  for(i in 1:B) {
    z=rnorm(n)
    L1=z %*% X
    L2=L1 %*% t(X)
    Lb2[i]=sum(L2^2)/m^2
  }
  
  return(Lb2)
}

randomization_Lb4 <- function(X, B, seed){
  # X: scaled genotype
  # B: iteration time
  set.seed(seed)
  
  n = nrow(X)
  m = ncol(X)
  
  Lb4=array(0, dim=B)
  for(i in 1:B) {
    z=rnorm(n)
    L1=z %*% X
    L2=L1 %*% t(X)
    L3=L2 %*% X
    L4=L3 %*% t(X)
    Lb4[i]=sum(L4^2)/m^4
  }
  
  return(Lb4)
}

randomization_Ly1 <- function(X, y){
  # X: scaled genotype (n*m)
  # y: phenotype (n*1)
  
  n = nrow(X)
  m = ncol(X)
  L1 = t(y) %*% X
  Ly1 = sum(L1^2)/m

  return(Ly1)
}

randomization_Ly2 <- function(X, y){
  # X: scaled genotype (n*m)
  # y: phenotype (n*1)
  
  n = nrow(X)
  m = ncol(X)
  L2 = t(y) %*% X %*% t(X)
  Ly2 = sum(L2^2)/m^2
  
  return(Ly2)
}

randomization_Ly3 <- function(X, y){
  # X: scaled genotype (n*m)
  # y: phenotype (n*1)
  
  n = nrow(X)
  m = ncol(X)
  L3 = t(y) %*% X %*% t(X) %*% X
  Ly3 = sum(L3^2)/m^3
  
  return(Ly3)
}


GenerateGeno <- function(freq, N) # return unrelated g( N*M )
{
  M = length(freq)
  g = matrix(NA, N, M)
  if(N==0) return(g)
  for(i in 1:M)
  {
    g[,i] = rbinom(N, 2, freq[i])
  }
  return(g)
}
```
